# Collagen I weakly interacts with the β-sheets of β_2_-microglobulin and enhances conformational exchange to induce amyloid formation

**DOI:** 10.1101/801522

**Authors:** Cody L. Hoop, Jie Zhu, Shibani Bhattacharya, Caitlyn A. Tobita, Sheena E. Radford, Jean Baum

## Abstract

Amyloidogenesis is significant in both protein function and pathology. Amyloid formation of folded, globular proteins is commonly initiated by partial unfolding. However, how this unfolding event is triggered for proteins that are otherwise stable in their native environments is not well understood. The accumulation of the immunoglobulin protein β_2_-microglobulin (β_2_m) into amyloid plaques in the joints of long-term hemodialysis patients is the hallmark of Dialysis Related Amyloidosis (DRA). While β_2_m does not form amyloid unassisted near neutral pH *in vitro*, the localization of β_2_m deposits to joint spaces suggests a role for the local extracellular matrix (ECM) proteins, specifically collagens, in promoting amyloid formation. Indeed, collagen and other ECM components have been observed to facilitate β_2_m amyloid formation, but the large size and anisotropy of the complex, combined with the low affinity of these interactions, has limited atomic-level elucidation of the amyloid-promoting mechanism by these molecules. Using solution NMR approaches that uniquely probe weak interactions and large complexes, we are able to derive binding interfaces for collagen I on β_2_m and detect collagen I-induced µs–ms timescale dynamics in the β_2_m backbone. By combining solution NMR relaxation methods and ^15^N-dark state exchange saturation transfer experiments, we propose a model in which weak, multimodal collagen I–β_2_m interactions promote exchange with a minor population of an amyloid-competent species to induce fibrillogenesis. The results portray the intimate role of the environment in switching an innocuous protein into an amyloid-competent state, rationalizing the localization of amyloid deposits in DRA.

## INTRODUCTION

Several proteins self-associate into amyloid fibrils, which in some cases have functional roles^1-3^, but for others are associated with debilitating human diseases^4-6^ including Alzheimer’s Disease, Parkinson’s Disease, Huntington’s Disease, type II diabetes, cataracts, and Dialysis Related Amyloidosis (DRA). The protein precursors of amyloid diseases have unrelated primary sequences and structures^7^, spanning natively unfolded (intrinsically disordered) states, such as α-synuclein, amyloid-β peptide, and tau^8-10^, to stable, globular proteins, such as β_2_-microglobulin (β_2_m), transthyretin, and immunoglobulin light chains^11-13^. Initiation of amyloid formation of the latter class of proteins requires unfolding or partial unfolding of monomeric precursors, which can transiently assume amyloid-competent state(s). This kinetic barrier may be lower for intrinsically disordered proteins. However, what triggers the initial unfolding and subsequent amyloidogenesis of natively folded globular proteins remains poorly understood.

Accumulation of β_2_m amyloid plaques in the joints of long-term hemodialysis patients leads to DRA and arthritic symptoms^14-17^. In healthy individuals, β_2_m dissociates from the major histocompatibility complex-I (MHC-I), is released into the plasma, and is carried to the kidneys for degradation^18-19^. However, when hemodialysis or peritoneal dialysis are required due to kidney failure, β_2_m is not efficiently removed from the plasma, leading to increased concentrations by up to 60-fold^16, 20-21^. Remarkably, despite being transported throughout the body, β_2_m accumulates into amyloid plaques specifically in skeletal tissues of dialysis patients^16, 21-24^. The mechanism(s) by which β_2_m fibrillizes *in vivo* is not well understood, since in isolation the wild-type protein (the major culprit of DRA) resists amyloid formation in physiological conditions, even at high (100 µM) concentrations^25-26^. It has been proposed that β_2_m amyloid localized in the joints could result, at least in part, from interactions with the major components of the extracellular matrix (ECM) in bone and cartilage: collagens I and II^20-23^ and glycosaminoglycans (GAGs)^27-29^. The binding affinities of β_2_m to these collagens have been shown to be in the µM–mM range^30^ with preference for collagen I^27^. Although the interaction is weak, it is nonetheless pathologically significant, as images of *ex vivo* DRA plaques reveal β_2_m amyloid covering the surface of collagen I fibrils^21^. Indeed, recent kinetics studies have revealed that ECM components, such as collagens^21, 28-29^ and GAGs^28-29, 31-32^, as well as pre-formed fibril seeds and other co-factors^25-26, 28, 31-49^, induce and modulate β_2_m amyloid formation. However, atomic details of how these components interact with, and induce, the amyloid formation of β_2_m have remained an open question.

The weak nature of the interaction and large, anisotropic shape of the β_2_m–collagen I complex create a challenge for deriving atomic-level information on how collagen I–β_2_m interactions initiate β_2_m amyloidogenesis. The immunoglobulin fold of monomeric β_2_m has dimensions of ∼ 4 nm × 2 nm × 2 nm, whereas the simplest triple helical unit of collagen I has strikingly larger dimensions of 300 nm × 1.5 nm × 1.5 nm. Collagen I triple helices assemble into even larger, structured fibrils that have diameters ranging from 10–500 nm and lengths on the µm-scale. Collagen I therefore presents as a large surface with numerous reactive groups for β_2_m interactions. These challenges are not insurmountable, however, as powerful solution nuclear magnetic resonance (NMR) spectroscopy methods can indirectly probe large, lowly populated complexes in site-specific detail that are invisible by other biophysical techniques.

In this study, by utilizing NMR spectroscopy experiments designed to probe large complexes, we are able to pinpoint the binding interface of wild-type β_2_m for collagen I at physiological pH and have shown it to involve both β-sheets of the native protein, suggestive of different binding modes for the same molecular complex. Residues identified at the binding interface include both hydrophobic and hydrophilic sidechains. Through ^15^N relaxation experiments, we have also found that collagen I increases the number of residues in β_2_m involved in conformational exchange on the µs–ms timescale. These regions include residues 6–11 (β-strand A), 36–39 (β-strand C), 51 (β-strand D), and 91–94 (β-strand G) in the edge β-strands and loop residues 15–20 (loop AB), 35 (loop BC), 52-53 (loop DE), 63 (loop DE), and 78 (loop EF), the dynamics and conformations of which are known to be important for β_2_m amyloid formation^31, 38, 50-51^. We propose that the weak interactions of collagen I with the β_2_m β-sheets promote exchange of the native protein with a minor population of amyloid-competent species that induce fibrillogenesis. This study illuminates how a protein component, collagen I, local to the environment in which β_2_m plaques are found, can interact with a stable, globular protein to initiate debilitating amyloid formation.

## RESULTS

### Collagen I induces β_2_m amyloid formation in a concentration-dependent manner

Since the direct interaction of β_2_m with collagen in the joint space has been proposed to induce β_2_m amyloid formation^21, 27^, we probed the β_2_m–collagen I interaction under physiological pH conditions (pH 7.4) using a solid-phase enzyme-linked immunosorbent assays (ELISA) (Figure 1A). Importantly, the results suggest a dose-dependent interaction of the two proteins, consistent with previously published results^27^, under the conditions employed here. The adhesion of β_2_m to casein was monitored as a negative control, for which no significant binding was observed (Figure 1A).

**Figure 1.**
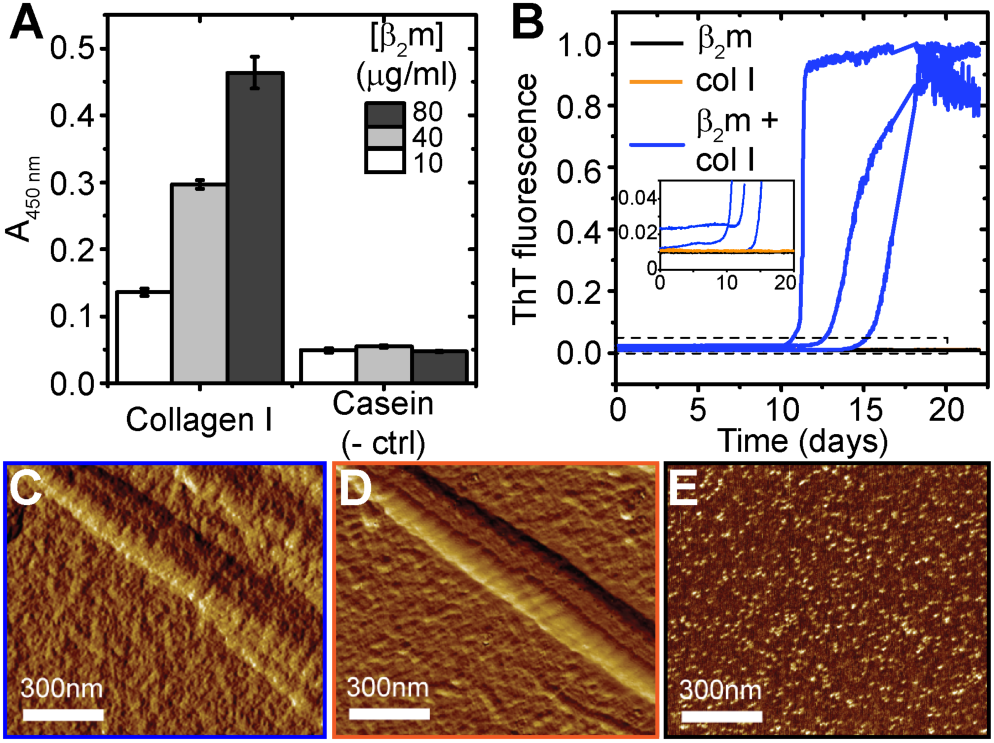
Detection of collagen I-driven β_2_m amyloid formation. (A) ELISA probing dose-dependent adhesion of β_2_m (10–80 µg/ml) to collagen I (10 µg/ml) or casein (10 µg/ml, used as a negative control), at pH 7.4. The average absorbance at 450 nm from triplicates within the same plate are reported with the standard deviation given as error bars. (B) ThT fluorescence curves of 85 µM β_2_m (black), 85 µM β_2_m + 3.4 mg/ml (8.5 µM) collagen I (blue), or 3.4 mg/ml collagen I alone (orange) over 22 days in 10 mM sodium phosphate buffer, pH 7.4, shaking at 600 rpm at 37°C. Three representative curves are given for each condition. The insert shows a zoom-in of the baseline of the ThT fluorescence curves to highlight the lack of fluorescence enhancement for both β_2_m (black) and collagen I (orange). (C–E) Representative amplitude-modulated AFM images of (C) β_2_m co-incubated with collagen I fibrils, (D) collagen I fibrils alone, and (E) β_2_m alone after incubating for four days at 37°C with shaking.

Having verified the β_2_m–collagen I interaction under physiological pH conditions, we next monitored amyloid growth of β_2_m in the presence or absence of collagen I by thioflavin T (ThT) fluorescence (Figure 1B). In the presence of 3.4 mg/ml collagen I (1:0.1 molar ratio β_2_m:collagen I), β_2_m amyloid is formed within 12–21 days (Figure 1B, blue), as evident by enhanced ThT fluorescence. This is not observed in the absence of collagen I in the same conditions, and collagen I alone does not show ThT fluorescence enhancement (Figure 1B). Notably, at lower concentrations of collagen I, lower β_2_m concentrations, or shorter timescales fibrils are not observed^29, 31^. Atomic force microscopy (AFM) images also show that β_2_m interacts with collagen I fibrils, with β_2_m coating the collagen I fibril surface before detectable fibril formation by ThT fluorescence, obscuring the characteristic collagen I fibril D-banding that is clearly observed in the absence of β_2_m (Figure 1C,D), consistent with previous results^21^. β_2_m alone (Figure 1E) does not aggregate in the conditions employed, with no fibrils or high molecular weight assemblies observed by AFM. These data confirm that adhesion of collagen I to β_2_m induces β_2_m amyloid formation under physiological conditions *in vitro*, while the protein is not able to form amyloid in the absence of collagen I.

### Weak, but specific β_2_m–collagen I interactions observed through ^15^N-R_2_ perturbations

In order to understand mechanistic details by which collagen I interacts with β_2_m to initiate amyloid formation, we used solution NMR methods, which provide an excellent toolbox of approaches able to characterize residue-specific features of weak protein–protein interactions on multiple timescales^52-55^. A titration of collagen I into a β_2_m monomer solution showed no significant chemical shift perturbations in ^1^H– ^15^N heteronuclear single quantum correlation (HSQC) spectra (Figure S1). However, a residue-specific attrition of the peak intensities observed with increasing collagen I concentrations (Figure 2A), suggests chemical exchange between the bound and free states of β_2_m consistent with the low affinity of the interaction in these conditions (K_d_ ≈ 410 µM^30^). To minimize collagen I aggregation during the NMR experiments and to capture the most specific interactions, we proceeded with low collagen I concentrations (0.6–1.2 mg/ml) that displayed consistent residue-specific perturbations and kept samples at 10°C, allowing NMR spectra to be acquired for over one week without visible alterations in spectral quality. Addition of 1.2 mg/ml collagen I to 300 µM β_2_m resulted in a reduction in resonance intensity of all peaks, consistent with transient formation of a high molecular weight complex (Figure 2A). However, the greatest reduction in peak intensities occurred for residues in the eight β-strands of the wild-type protein (Figure 2A). These peak intensity losses are in part due to increased ^15^N-transverse relaxation rates (R_2_), which are sensitive to changes in internal motions on the ps–ns timescale and conformational exchange on the µs–ms timescale. Indeed, at these concentrations, we observe an overall increase in ^15^N-R_2_, but importantly, the increase is not uniform across all residues, but is residue specific, involving predominantly residues 2–3 (N-terminus), 7–11 (β-strand A), 16–19 (loop AB), 23–26 (β-strand B), 35–39 (β-strand C), 50–52 (β-strand D), 64, 66–69 (β-strand E), 79–82 (β-strand F), 85, 87 (loop FG), and 91–94 (C-terminal β-strand G) (Figure 2B-C). The increased ^15^N-R_2_ at these specific sites could have multiple origins, arising due to reduced backbone mobility upon direct interaction with collagen I and/or to line broadening due to exchange between species with different chemical shifts, especially since the observed ^15^N-ΔR_2_ is dependent on magnetic field (700 MHz vs. 900 MHz, Fig. S2). In order to disentangle these contributions to the increase in ^15^N-R_2_, we proceeded with two sets of NMR experiments: ^15^N-dark state exchange saturation transfer (DEST) experiments, which can identify residues interacting with the large complex, and in-phase Hahn-echo experiments, which detect conformational exchange on the µs–ms timescale.

**Figure 2.**
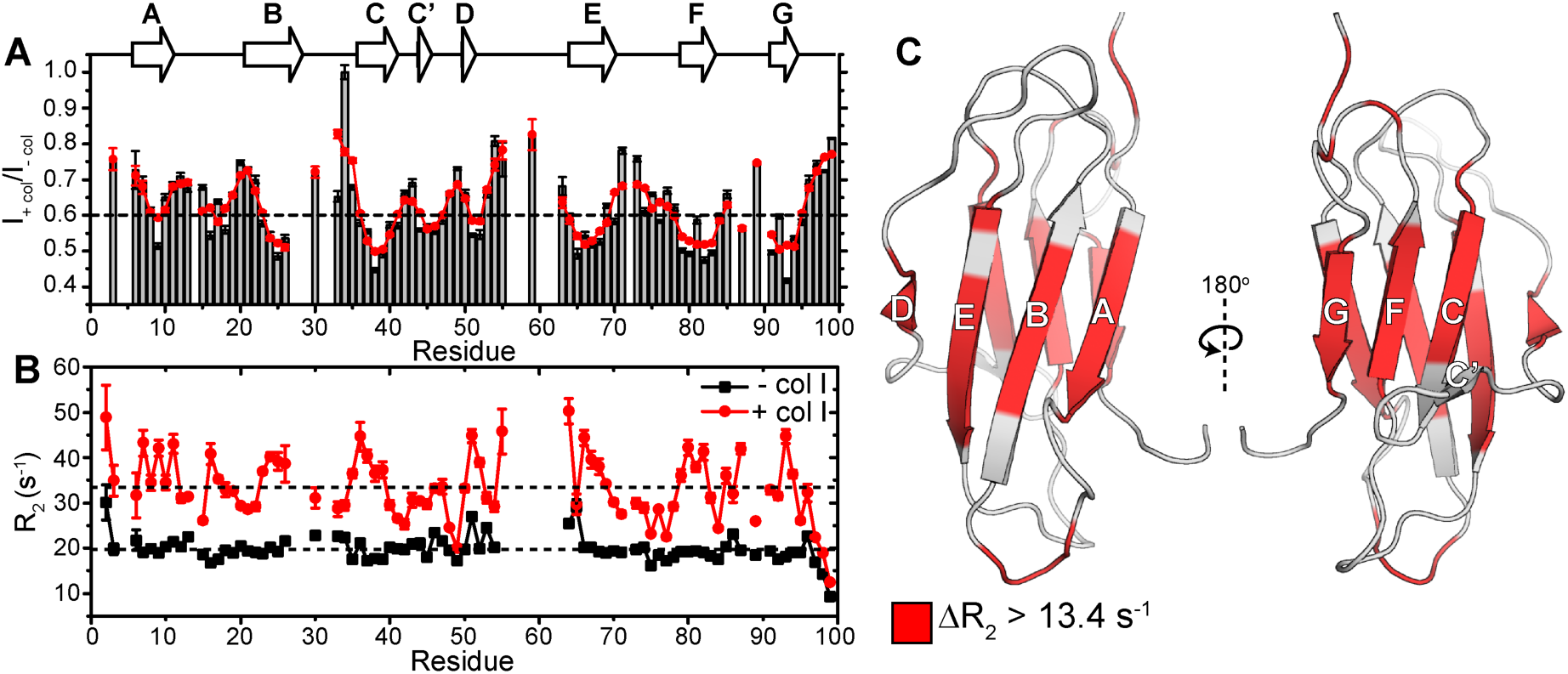
Characterizing residue-specific β_2_m–collagen I binding through ^15^N-R_2_ measurements. (A) Amide backbone signal intensity ratios from ^1^H–^15^N HSQC spectra of 300 µM β_2_m in the presence of 1.2 mg/ml collagen I compared with in the absence of collagen I (gray bars). The red line is a smoothened curve of the signal intensity ratios to help guide the eye. The dashed line is drawn at the average signal intensity ratio over the entire protein. Dips in the signal intensity reflect regions maximally perturbed by the presence of collagen I. Error bars are propagated from the noise level of the spectra. The secondary structure is indicated above the plot. (B) ^15^N-R_2_ measurements of 300 µM β_2_m in the presence (red) or absence (black) of 1.2 mg/ml collagen I. The errors are propagated from the fitting errors. The dashed lines indicate the mean ^15^N-R_2_ values of β_2_m in the presence or absence of 1.2 mg/ml collagen I over the entire protein. All experiments were conducted in TBS, pH 7.4 containing 0.5 mg/ml casein as a non-specific binding blocking agent at 10°C. Note that in these conditions, several residues in the DE loop do not have observable peak intensities in the ^1^H–^15^N HSQC spectrum due to inherent conformational exchange, consistent with previous results^49^. (C) Solution NMR structure of the WT-β_2_m monomer (PDB: 2XKS^49^) highlighting residues that show an increase in ^15^N-R_2_ higher than 13.4 s^-1^ (the mean Δ^15^N-R_2_) upon addition of 1.2 mg/ml collagen I.

### Pinpointing the collagen I interaction interface on β_2_m through ^15^N-DEST

In order to determine which residues of β_2_m interact most intimately with collagen I, we used ^15^N-DEST experiments^56-57^. This experiment is optimal when there is a measurable increase in R_2_ due to formation of a transient, large complex that is NMR-invisible because of its high R_2_ and detects the exchange between an observable ‘light’ state (free monomeric β_2_m) and the NMR-invisible ‘dark’ state (the high molecular weight collagen I–β_2_m complex). In the DEST experiment, high molecular weight species with high R_2_ values, such as the collagen I–β_2_m complex, can be partially saturated by weak radiofrequency (RF) fields at frequency offsets where monomeric β_2_m is not saturated. Saturation transfer to the observable monomeric species by chemical exchange is detected as a loss in monomeric β_2_m signal intensity. The broadening of these DEST saturation profiles (reduced signal intensities at further frequency offsets) in the presence of collagen I, relative to in its absence, is therefore indicative of residues at the interaction interface (Figure 3A–B). The ‘broadness’ of the profiles was measured by calculating the DEST difference (Θ) for each residue, which is a measure of the relative effects of on-resonance and off-resonance ^15^N saturation. Using a saturation frequency of 350 Hz, we measured Θ as: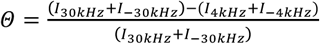, where ±30 kHz were the most off-resonance ^15^N offsets, and ^15^N offsets of ±4 kHz provide enough saturation transfer from bound to unbound β_2_m to show significant intensity loss without eliminating the signal in most cases. A substantial change in Θ (ΔΘ) upon addition of collagen I is reflective of residues at the binding interface (Figure 3A). Notably, we observe that the broadening of the DEST saturation profiles is residue-specific and not uniform across all β_2_m residues, with some residues showing no change in the DEST difference in the presence of collagen I (Figure 3A–B). Examples of DEST profiles in the presence or absence of collagen I for a residue that shows DEST due to collagen I binding (V82 in β-strand F) and one that does not (K41 in the C–C’ loop) are given in Figure 3B. In Figure 3A, those residues with ΔΘ larger than the mean, and likely have the most direct contacts with the collagen in the β_2_m–collagen I complex (shaded red), include residues 6–11 (β-strand A), 15–20 (loop AB), 21–26 (β-strand B), 35 (loop BC), 36–39 (β-strand C), 51 (β-strand D), 52–53 (loop DE), 63 (loop DE), 64–69 (β-strand E), 78 (loop EF), 79–83 (β-strand F), and 91–94 (β-strand G).

**Figure 3.**
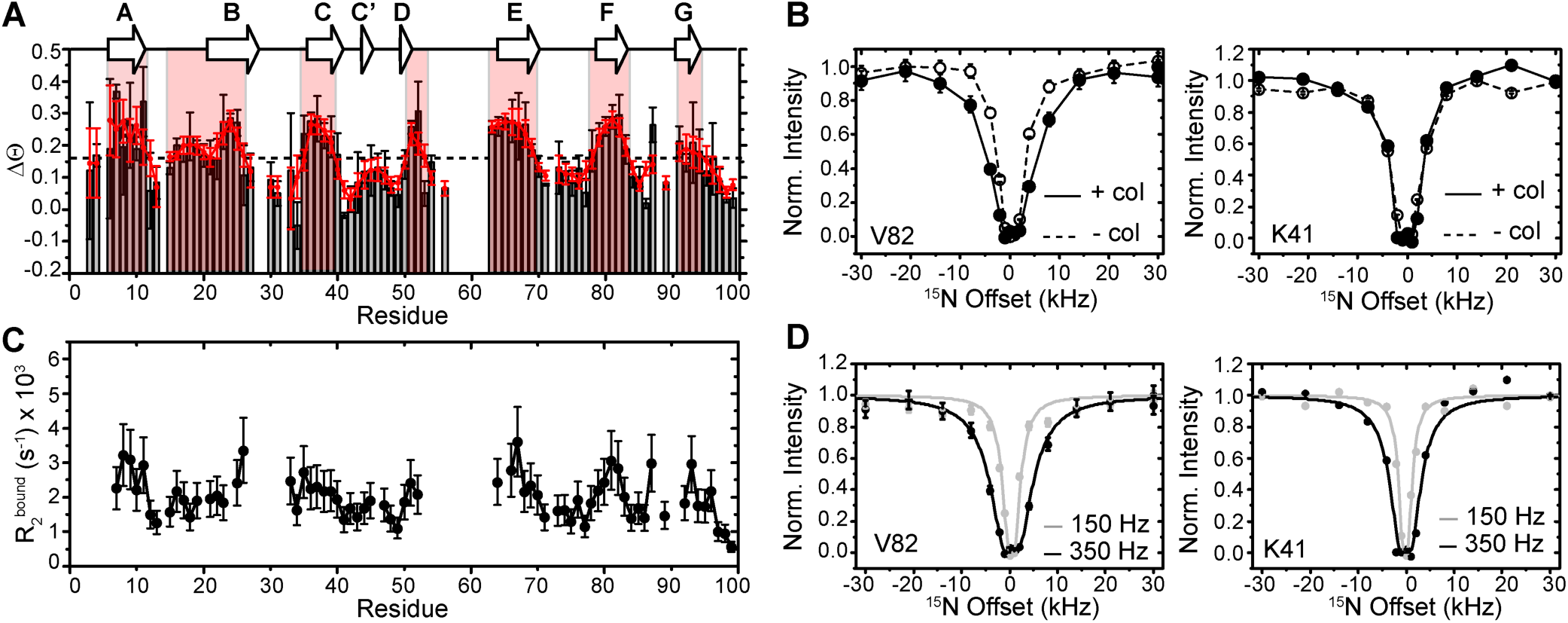
^15^N-DEST to identify collagen-binding interface on ^15^N-β_2_m. (A) ΔΘ calculated from ^15^N-β_2_m DEST intensities at ±30 kHz and ±4kHz 15N offsets with a 350 Hz saturation frequency in the presence or absence of collagen I (gray bars). The red line is a smoothened curve to guide the eye. The dashed line represents the average value of ΔΘ from all residues. The secondary structure is shown above the plot. (B) Examples of ^15^N-DEST profiles of 300 µM ^15^N-β_2_m in the presence (solid line, solid circles) or absence (dashed line, open circles) of 0.6 mg/ml collagen I. V82 in β-strand F shows enhancement of the DEST effect upon addition of collagen, whereas K41 in the CC’ loop does not. (C) ^15^N-R_2_^bound^ for each residue determined from fitting ^15^N-DEST profiles to the McConnell equations. (D) Examples of fit DEST profiles using the same residues as in (B). Data points are shown as circles and the fits as solid lines (gray-150 Hz saturation, black-350 Hz saturation). All experiments were carried out in TBS, pH 7.4, 10°C at 700 MHz ^1^H Larmor frequency.

In addition, the full DEST profiles can be used to quantify residue-specific transverse relaxation values of β_2_m in the collagen I-bound state (R_2_^bound^) and exchange kinetics between the bound and unbound β_2_m. Since the ΔR_2_ may be due to more complex processes than collagen I binding alone, such as an overall increased viscosity due to the presence of the large collagen I molecules, we fit only the ^15^N DEST profiles of each residue with 150 Hz and 350 Hz RF saturation to the McConnell equations^56-57^. Fitting to a simple two-state model, the population of the unbound, monomeric β_2_m was determined to be 94 +/- 2% with an apparent first-order rate constant for the conversion of β_2_m from unbound to collagen I-bound conformation (k_on_^app^) of 6.4 +/- 0.8 s^-1^. We interpret the direct binding interface to be the residues with the highest R_2_^bound^. The ^15^N-R_2_^bound^ profile shows a similar trend to the ΔΘ profile (Figure 3A, C), and suggests that binding interfaces for collagen I on β_2_m occur on both β-sheets. Examples of fitting to the experimental values of residues V82 (in a binding region) and K41 (away from interface) are shown in Figure 3D.

### Collagen I induced conformational exchange in β_2_m revealed by ^15^N relaxation

The enhanced ^15^N-R_2_ of β_2_m may not only be due to binding with a high molecular weight species (such as in a large complex), but also to an increase in conformational exchange dynamics of β_2_m on the µs–ms timescale, since the ^15^N-ΔR_2_ is dependent on the magnetic field (Figure S2). In order to determine which residues in β_2_m are in conformational exchange in the presence of collagen I, we use ^15^N in-phase Hahn echo experiments (R_2_^HE^) to estimate the relaxation exchange rates. At pH 7.4 and 10°C, few residues in β_2_m have R_ex_ values greater than 10s^-1^ in the absence of collagen I as measured by the in-phase Hahn echo experiments (Figure 4A). The N-terminus and residues in the BC and DE loops (for which several signals are unobservable) are natively in conformational exchange (Figure 4A). Upon addition of 0.6 mg/ml collagen I, the regions with high R_ex_ are expanded to include the full N-terminal β-strand A, part of β-strand B to part of β-strand C, including the connecting BC loop, β-strand D, the DE loop, the C-terminal end of β-strand F into the FG loop, and the C-terminal β-strand G (Figure 4B). Conformational dynamics in specific regions of β_2_m, including the N-terminal region and the BC loop that contains *cis* Pro32, have been shown to be crucial in controlling the amyloidogenicity of the protein^49, 58^. Thus, the enhanced conformational exchange induced by the presence of collagen I may facilitate minor populations of amyloid-component states of β_2_m.

**Figure 4.**
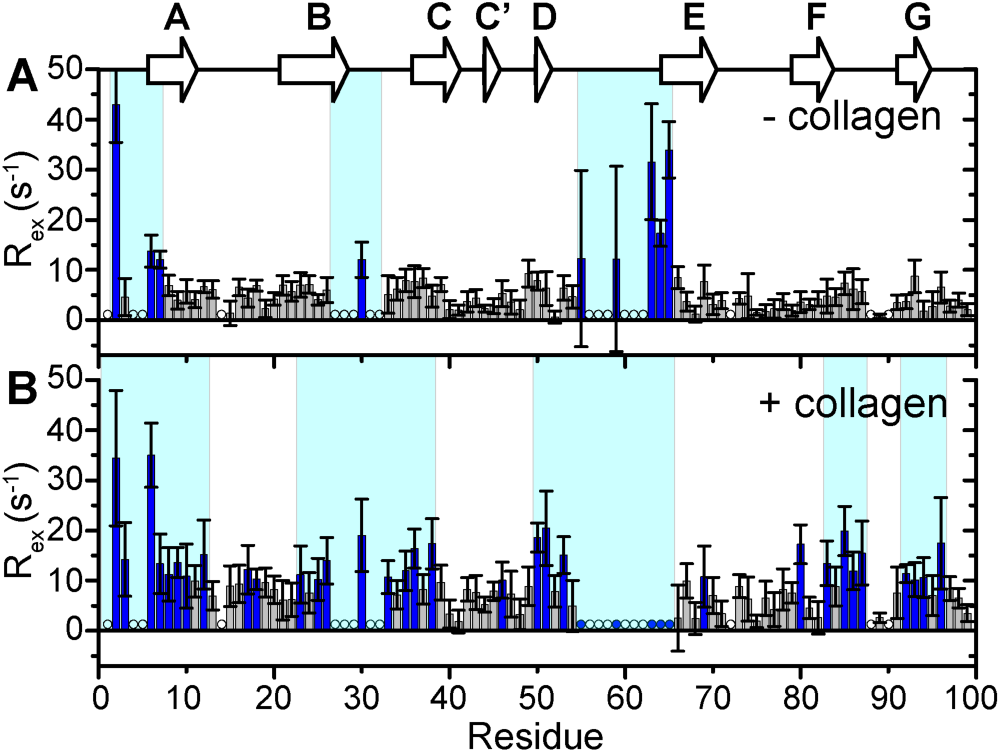
Conformational exchange in β_2_m induced by collagen I. Relaxation exchange rates (R_ex_) obtained by ^15^N-R_2_ Hahn echo experiments for each residue in 300 µM ^15^N-β_2_m in the absence (A) or presence (B) of 0.6 mg/ml collagen I at pH 7.4, 10°C, 700 MHz ^1^H Larmor frequency. R_ex_ values over 10 s^-1^ are indicated in blue bars. Regions shaded in cyan in both panels contain several residues with R_ex_ >10 s^-1^ in the respective conditions. Residues with unobservable cross-peaks are indicated by open circles. Residues that were observable in the absence of collagen I but were reduced to the level of the noise in the presence of collagen I are indicated by filled blue circles. Error bars are propagated from fitting errors.

## DISCUSSION

### A novel collagen I binding surface on β_2_m

Amyloid formation of β_2_m at physiological pH *in vitro* requires assistance by co-factors^21, 25-26, 28-29, 31-49^. In particular, ECM molecules, such as collagens and GAGs have been targeted as amyloid-inducing co-factors, since β_2_m amyloid formation has been localized to musculoskeletal tissues^16, 22-24^. While previous experiments have focused on the kinetics of amyloid formation in the presence of these molecules^21, 28-29, 31, 59^, a detailed atomistic description of the interactions involved and how these may enhance β_2_m conformational dynamics and amyloid formation had not been elucidated. Here, we have used complementary NMR relaxation-based experiments to pinpoint residues of β_2_m involved in the collagen I binding interface and collagen I-induced dynamics that lead to enhanced β_2_m amyloid formation at neutral pH *in vitro*. The ^15^N-DEST experiments indicate that residues in β-strands A, B, C, D, E, F, and G form interaction surfaces with collagen I. These provide two surfaces of mixed hydrophilic and hydrophobic composition (Fig. S3). Both contain hydrophobic patches with the ABED β-sheet displaying several aromatic residues on the interaction surface (Fig. S3). Since both β-sheets on opposite sides of the molecule were determined to interact with the collagen I surface, binding must be multimodal involving interaction surfaces formed by K6, Q8, Y10, F22, N24, Y26, S52, Y63, L65, Y67, and E69 on the ABED β-sheet and E36, D38, L40, A79, R81, N83, I92, and K94 on the GFC β-sheet (Figure 3, S3). Comparison of the molecular dimensions of the interacting molecules (4 x 2 x 2 nm for β_2_m, 300 nm x 1.5 nm for a collagen I triple helix, and microns in length x up to 500 nm in diameter for mature collagen I fibrils) highlights the potential for a myriad of binding modes, enabling independent binding of several β_2_m molecules to the same collagen molecule (Figure 5A–B). Importantly, the collagen I triple helix surface is interspersed with numerous hydrophilic and hydrophobic residues along its length (Figure 5A). The collagen I fibril surface maintains this repeating pattern of surface chemistries (Figure 5B), enhancing the potential for multiple binding modes to complementary surfaces in β_2_m (Figure 5C).

**Figure 5.**
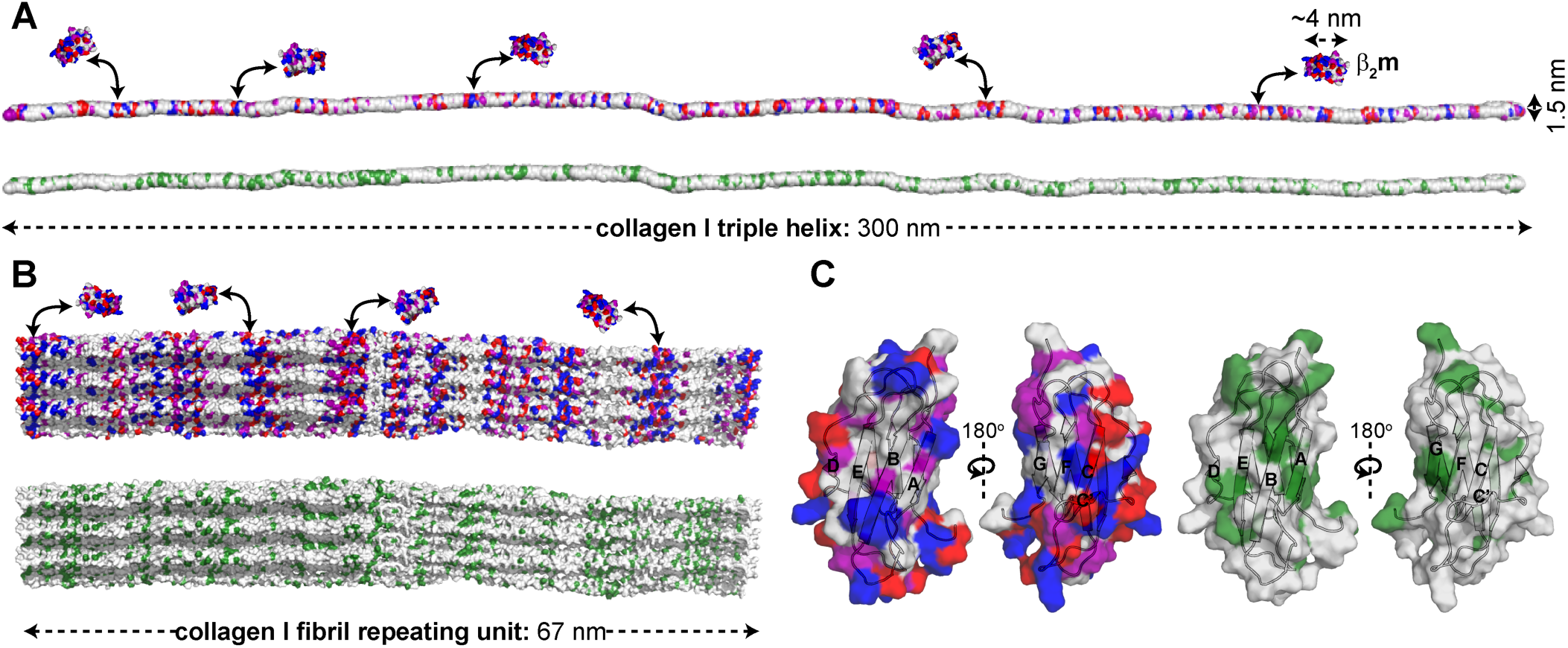
Potential surface contacts for β_2_m–collagen I interaction. A) Surface model of the collagen I monomer (PDB: 3HKS^60^) color-coded by amino acid type (top: hydrophilic, bottom: hydrophobic). A surface representation of β_2_m is shown for size comparison (PDB: 2XKS^49^). As an example, electrostatic surfaces of β_2_m are shown weakly interacting with electrostatic surfaces of collagen I. (B) Surface models of the collagen I fibril repeating unit (built from PDB: 3HKS^60^), color-coded by amino acid type (top: hydrophilic, bottom: hydrophobic). The repeating unit is ∼67 nm in length, however mature fibrils can be microns long and ∼500 nm in diameter. Distinct bands of electrostatic residues are observed within the repeating unit. (C) Surface representation of β_2_m monomer (PDB: 2XKS^49^) color-coded by amino acid type (left: hydrophilic, right: hydrophobic). All models are color-coded as: red, acidic; blue, basic; purple, uncharged-polar; and green, hydrophobic.

Collagen I is known to interact with multiple immunoglobulin-like protein folds through binding interfaces that include both hydrophobic and hydrophilic residues. Interactions of collagen I with osteoclast-associated receptor (OSCAR), leukocyte-associated immunoglobulin-like receptor-1 (LAIR-1), and glycoprotein VI (GPVI), play functional roles in immune system regulation^61-64^ and platelet activation^65-67^. Similar to the β-sheet binding interface on β_2_m for collagen I identified here, the collagen I binding sites on OSCAR and LAIR-1 are also found in β-sheet regions^68-69^. In the case of the OSCAR–collagen I interactions, Tyr and Arg residues that line the interacting β-sheet of OSCAR have been suggested to play a primary role^68^. LAIR-1 binds primarily to collagen fragments rich in Gly, Pro and hydroxyproline (GPO) content, but also has been shown to interact with multiple binding motifs in collagen II and III toolkit peptides, some of which are not GPO rich^69^. NMR and mutagenesis studies on LAIR-1 have shown that depletion of Arg or Glu at the putative β-sheet interface showed decreased collagen binding, suggesting a role for electrostatic interactions^70^. GPVI, also recognizes GPO rich collagen motifs, however through a unique hydrophobic groove formed by a β-strand connecting loop that is flanked by hydrophilic residues^71-73^. Thus, although these proteins all share a similar immunoglobulin fold, each shows a unique binding interface to collagen, interacting in grooves formed by β-sheets or loops and having both hydrophobic and hydrophilic residues that each play fundamental roles in binding.

### Collagen-induced conformational dynamics in β_2_m reflect amyloid prone dynamics

Beyond the structured collagen I-binding interface of β_2_m, using ^15^N relaxation experiments, we observe enhanced dynamics in the N- and C-termini, BC and FG loops, and the β-strand D of β_2_m upon complex formation. Enhanced dynamics in each of these regions has been proposed to play key roles in the aggregation mechanism of wild-type β_2_m^38, 49-51, 74-84^. Amyloid formation of β_2_m is nucleation dependent and proceeds through a near native folding intermediate, I_T_, that is in part defined by a non-native *trans*-His31-Pro32 peptide bond in the BC loop^38, 50-51, 85-86^. The *cis-trans* isomerization of Pro32 is aided by displacement of the N-terminal six residues, which destabilizes the BC loop, allowing β_2_m to sample multiple amyloidogenic conformations that enhance the rate of aggregation^38, 50-51, 74, 79-81, 85-86^. Deletion of the first six N-terminal residues in the naturally occurring variant, ΔN6, enhances the propensity for amyloid formation, and aggregation occurs in the absence of additional cofactors at physiological pH *in vitro*^49, 74, 87-88^. In addition, NMR relaxation experiments show enhanced dynamics in β-strand D and the DE loop of amyloidogenic ΔN6^49^, which have been proposed to contribute to its higher aggregation propensity. NMR studies of a P32G-β_2_m variant, which inherently has a *trans*-His31-Gly32 peptide bond, showed significant line broadening in β-strands A and D and the BC and FG loops relative to WT-β_2_m^38^. This was interpreted to result from conformational conversion between the native and I_T_ conformations^38^. The observation of increased R_ex_ of these same regions upon addition of collagen I to WT-β_2_m, in this study, is consistent with the same regions undergoing conformational exchange from the native state to an amyloidogenic precursor consistent with the I_T_ state, to enhance amyloid formation. Such a model provides a mechanism to enhance *cis-trans* Pro isomerization to initiate assembly into amyloid without the involvement of a prolyl isomerase.

### A proposed mechanism of collagen I-driven β_2_m amyloidogenesis

With the new insights into the binding interface of collagen I on β_2_m and its impact on β_2_m dynamics described here, we propose a mechanistic view of how collagen I drives amyloidogenesis of β_2_m. In the presence of collagen I, the β-sheets of β_2_m are available for binding to the collagen I surface, with both β-sheets providing potential binding interfaces, indicative of multiple binding modes, rather that a unique and specific binding interface. The interaction between the two molecules is mediated by hydrophobic and electrostatic interactions (Figures 3, S3). In its native state, high transverse relaxation rates are observed in the apical loops of β_2_m, including the BC loop that contains *cis* Pro32 and the adjacent DE loop (Figure 2B). Additional dynamics upon collagen I binding are imposed on the N-terminus, β-strands B and C, BC loop, β-strand D, FG loop, and the C-terminal β-strand G (Figure 4B). Through modification of the dynamics of β_2_m in these sites, the probability of *cis-trans* isomerization of Pro32, known to be a key step in β_2_m fibril formation^85-86^, will be increased, with concomitant sampling of amyloid-competent species, including the I_T_ state, known to promote amyloid formation^38, 50-51^ (Figure 6). The results provide a molecular explanation for the mechanism of deposition of β_2_m in collagenous-rich joints in dialysis patients^16, 21-24^. More generally, they also serve as an exemplar of the key role of the physiological environment in amyloid formation, by rationalizing the often remarkably specific deposition of amyloid to different tissues^1^, and in some cases, of different variants of the same protein in different tissues^89-90^. The methods used here to interrogate the weak-transient interaction of the large, β_2_m–collagen I complex can be extended to future studies to gain atomic-level insight into how other physiologically relevant cofactors promote amyloid formation of globular proteins involved in other amyloid diseases.

**Figure 6.**
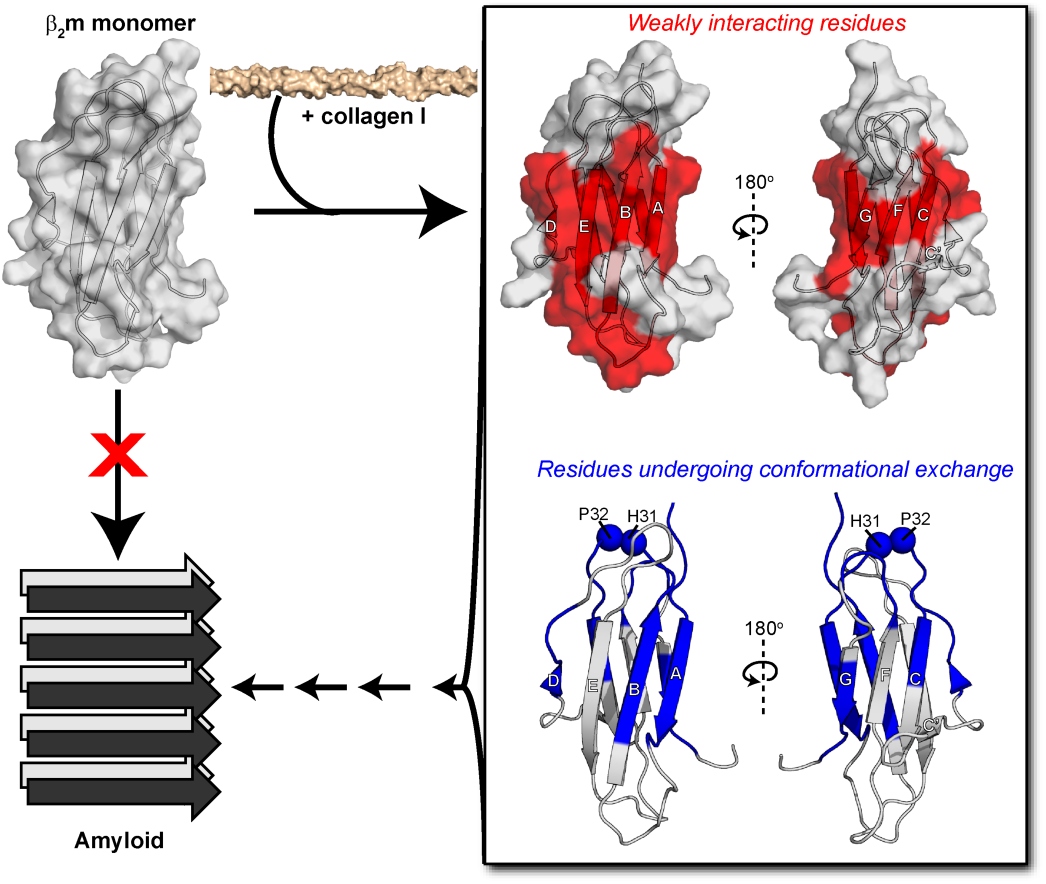
Proposed mechanism for collagen-driven β_2_m amyloidogenesis. Alone, the β_2_m monomer (PDB: 2XKS^49^) does not readily aggregate into amyloid fibrils. Upon addition of collagen I, we have observed an interaction interface to include both β-sheets of β_2_m through the ^15^N-DEST experiment (red). Collagen also induces conformational exchange in regions colored in blue, as assessed by ^15^N relaxation experiments. The interaction of collagen I with the structured regions of β_2_m enhances conformational exchange, promoting formation of an amyloid-competent species and inducing aggregation.

## MATERIALS AND METHODS

### Expression and purification of β_2_m

Wild-type β_2_m was expressed recombinantly in *Escherichia coli* BL21(DE3) pLysS cells by induction with 1 mM IPTG overnight at 37°C, following methods described previously^39^. Cells were lysed in 25 mM Tris-HCl buffer, pH 8.0 and with an Avestin Emulsiflex-C5 homogenizer. β_2_m is accumulated in inclusion bodies. To extract the β_2_m from inclusion bodies, the cell pellet was washed five times with 25 mM Tris-HCl buffer, pH 8.0 and solubilized in 25 mM Tris-HCl, pH 8.0 buffer containing 8 M urea, rocking overnight at room temperature. The protein was verified to be in the soluble fraction by SDS-PAGE. β_2_m was refolded by dialyzing against 25 mM Tris-HCl buffer, pH 8.0 at 4°C and purifying by anion exchange (HiTrap Q HP, GE Healthcare). The protein was further purified by size exclusion chromatography with a Superdex 75 gel filtration column (GE Life Sciences). Protein purity was verified by SDS-PAGE, and concentrations for experiments were determined by measuring the absorbance at 280 nm using a molar extinction coefficient of 19,060 M^-1^cm^-1^. [U-^15^N]-enriched β_2_m was expressed recombinantly for NMR using the same protocol in HCDM1 minimal media supplemented with ^15^N-ammonium chloride.

### ELISA

Relative adhesion of variable concentrations of β_2_m to collagen I was determined by ELISA experiments. Nunc Maxisorp 96-well plates (Thermo Scientific) were coated with 100 µl of collagen I from rat tail tendon (BD Biosciences; 10 µg/ml in 10 mM acetic acid) overnight at 4°C. Uncoated areas on the plates were blocked with 200 µl of 0.5% w/v casein in binding buffer at room temperature for 1 hr. The binding and washing buffer consisted of PBS at pH 7.4 with 0.05% v/v Tween 20 (PBS-T) and 0.05% w/v casein as a non-specific blocking agent. After washing the wells three times with 200 µl washing buffer, 100 µl β_2_m in PBS-T and 0.05% w/v casein (10 µg/ml, 40 µg/ml, or 80 µg/ml) was added to the wells and incubated for 1 hr at room temperature. After three washes with 200 µl washing buffer, 100 µl mouse anti-β_2_m monoclonal antibody (1:2000 v/v in PBS-T and 0.05% w/v casein, Millipore Sigma) was bound to β_2_m in each well by incubating at room temperature for 1 hr. Subsequently, following three washes with 200 µl washing buffer, 100 µl of goat HRP-conjugated anti-mouse secondary antibody (1:5000 v/v dilution in PBS-T and 0.05% w/v casein, Genscript) was incubated in the wells at room temperature for 30 min. After washing for a final four times with 200 µl washing buffer, the binding of β_2_m to collagen I was detected through a colorimetric assay using a 3,3’,5,5’-tetramethylbenzidine substrate kit (Pierce) according to the manufacturer’s protocol, and measuring the absorbance at 450 nm using a Tecan Infinite F50 plate reader with Magellan software.

### ThT fluorescence

Amyloid fibril formation was monitored by ThT fluorescence assays of β_2_m in the presence or absence of collagen I fibrils. Purified recombinant β_2_m lyophilized powder was dissolved in 100 µl of 10 mM sodium phosphate buffer, pH 7.4 to 1 mg/ml (85 µM). Collagen I fibrils were prepared by incubating 3.4 mg/ml collagen I (BD Biosciences) in PBS, pH 7.4 at 37°C for 1 hr. The fibril suspension was sonicated in a bath sonicator for 10 min and centrifuged at 16,500 rpm for 10 min to isolate fibrils. Collagen fibril pellets were resuspended in 100 µl of 10 mM sodium phosphate buffer, pH 7.4 in the presence or absence of β_2_m. Three or four samples were prepared for each condition and were transferred to a 96-well plate. ThT was added to each sample to a final concentration of 10 µM. ThT fluorescence was monitored over 22 days at 37°C with shaking at 600 rpm in a POLARstar Omega fluorimeter (BMG Labtech).

### NMR

For all NMR experiments, purified recombinant [U-^15^N]-labeled β_2_m was diluted to 300 µM in TBS, pH 7.4 with 0.5 mg/ml casein and 10% v/v D_2_O. Before mixing, collagen I from rat tail tendon was dialyzed against TBS, pH 7.4. The concentration of collagen I after dialysis was determined by bicinchoninic acid assay (Pierce). All experiments were performed at 10°C. All data were collected on a 700 MHz Bruker *AVIII* or 900 MHz *AVI* NMR spectrometers equipped with TCI-cryo-probes. Data were processed in NMRPipe^91^ and analyzed in Sparky^92^.

#### ^1^H-^15^N HSQC spectra

^1^H-^15^N HSQC spectra^93-94^ of [U-^15^N]-labeled β_2_m were acquired with different concentrations of collagen I (0, 0.12 mg/ml, and 1.2 mg/ml) in TBS, pH 7.4 with 0.5 mg/ml casein and 10% D_2_O at 10°C. The intensity ratio is taken as the intensity of a given cross-peak in the ^1^H-^15^N HSQC spectrum of β_2_m in the presence of collagen I relative to the intensity of the same cross-peak in the absence of collagen I, determined in Sparky^92^. The errors were propagated from the signal to noise ratio in each spectra.

#### ^15^N-R_2_ and ^15^N-R_2_^HE^

[U-^15^N]-labeled β_2_m ^15^N transverse relaxation rates (R_2_) were measured from a series of HSQC-based 2D ^1^H-^15^N spectra using the Carr-Purcell-Meiboom-Gill (CPMG) pulse sequence^95^ with varying relaxation delays: in the absence of collagen I at 700 MHz-0, 16, 16, 32, 48, 64, 64, 80, 96, and 112 ms and 900 MHz-0, 16, 32, 32, 32, 48, 64, 80, 96, 112, and 128 ms and in the presence of 1.2 mg/ml collagen I at 700 mHz-0, 16, 16, 32, 48, 48, 64, and 80 ms and at 900 MHz-0, 16, 32, 32, 32, 48, 64, 80, and 96 ms. Relaxation delays used to quantify ^15^N-R_2_ rates of β_2_m in the presence of 0.6 mg/ml collagen I at 700 MHz were: 0, 8, 8, 16, 24, 32, and 56 ms and in the absence of collagen I: 0, 8, 8, 16, 24, 40, and 56 ms. ^15^N-R_2_ ^HE^ informs on the chemical exchange contribution to R_2_ by using an in-phase Hahn echo experiment^96^. Relaxation delays used in the R_2_^HE^ experiment both in the presence and absence of 0.6 mg/ml collagen I were: 0.768, 7.68, 7.68, 15.4, 23, 38.5, and 61.4 ms. In each case, the R_2_ rates were determined by fitting peak intensities to a single exponential decay function. The chemical exchange contribution (R_ex_) for each β_2_m residue in the absence and presence of 0.6 mg/ml collagen I was determined as: R_ex_ = R_2_^HE^-R_2_.

#### DEST experiments

The ^15^N-DEST experiment^56-57^ was applied to [U-^15^N]-labeled β_2_m in the presence or absence of 0.6 mg/ml collagen I. In this experiment, an ^15^N saturation pulse of 150 or 350 Hz was applied for 0.9 ms at different ^15^N frequency offsets: 0, ±1, ±2, ±4, ±8, ±14, ±21, and ±30 kHz. An experiment in which the ^15^N saturation pulse was set to 0 Hz with an offset of 30 kHz was also included as a reference. The ^15^N-DEST profiles were extracted for each residue as the peak intensity at each ^15^N saturation offset and were fitted to a two-state model using the destfit program by Clore and co-workers to obtain R_2_^bound^, p_bound_, and k_on_^app56-57^. The ΔΘ profile was obtained by measuring Θ for each β_2_m residue in the presence and absence of 0.6 mg/ml collagen I as: 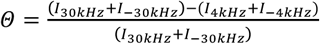, and taking ΔΘ = Θ_+col_ − Θ_−col_

## ASSOCIATED CONTENT

### Supporting Information

The Supporting Information includes figures that show an overlay of ^1^H-^15^N HSQC spectra of [U-^15^N]-β_2_m in the presence of 0 mg/ml, 0.12 mg/ml, or 1.2 mg/ml collagen I (Figure S1); ^15^N-R_2_ measurements of [U-^15^N]-β_2_m in the absence or presence of collagen I at 700 MHz or 900 MHz (Figure S2); and a model showing the amino acid composition of the interacting β_2_m β-sheets (Figure S3). (PDF)

## AUTHOR INFORMATION

### Notes

The authors declare no competing financial interests.

## ACKNOWLEDGMENT

We acknowledge Arthur Palmer for helpful discussions. We also acknowledge with thanks the many discussions with our group members. We thank Ana Monica Nunes for contributions in the beginning of this project and Nuria Benseny-Cases, who provided critical insights in the early stages of the work.

## ABBREVIATIONS

β_2_m: β_2_-microglobulin;
DRA: Dialysis Related Amyloidosis;
ECM: extracellular matrix;
MHC-I: major histocompatibility complex-I;
GAG: glycosaminoglycan;
NMR: nuclear magnetic resonance;
ELISA: enzyme-linked immunosorbent assay;
ThT: thioflavin T;
AFM: atomic force microscopy;
HSQC: heteronuclear single quantum correlation;
DEST: dark-state exchange saturation transfer;
OSCAR: osteoclast associated receptor;
LAIR-1: leukocyte associated immunoglobulin-like receptor-1;
GPVI: glycoprotein VI;
CPMG: Carr-Purcell-Meiboom-Gill.

