## Supporting Information for "Collagen I weakly interacts with the β-sheets of β_2_-microglobulin and enhances conformational exchange to induce amyloid formation"

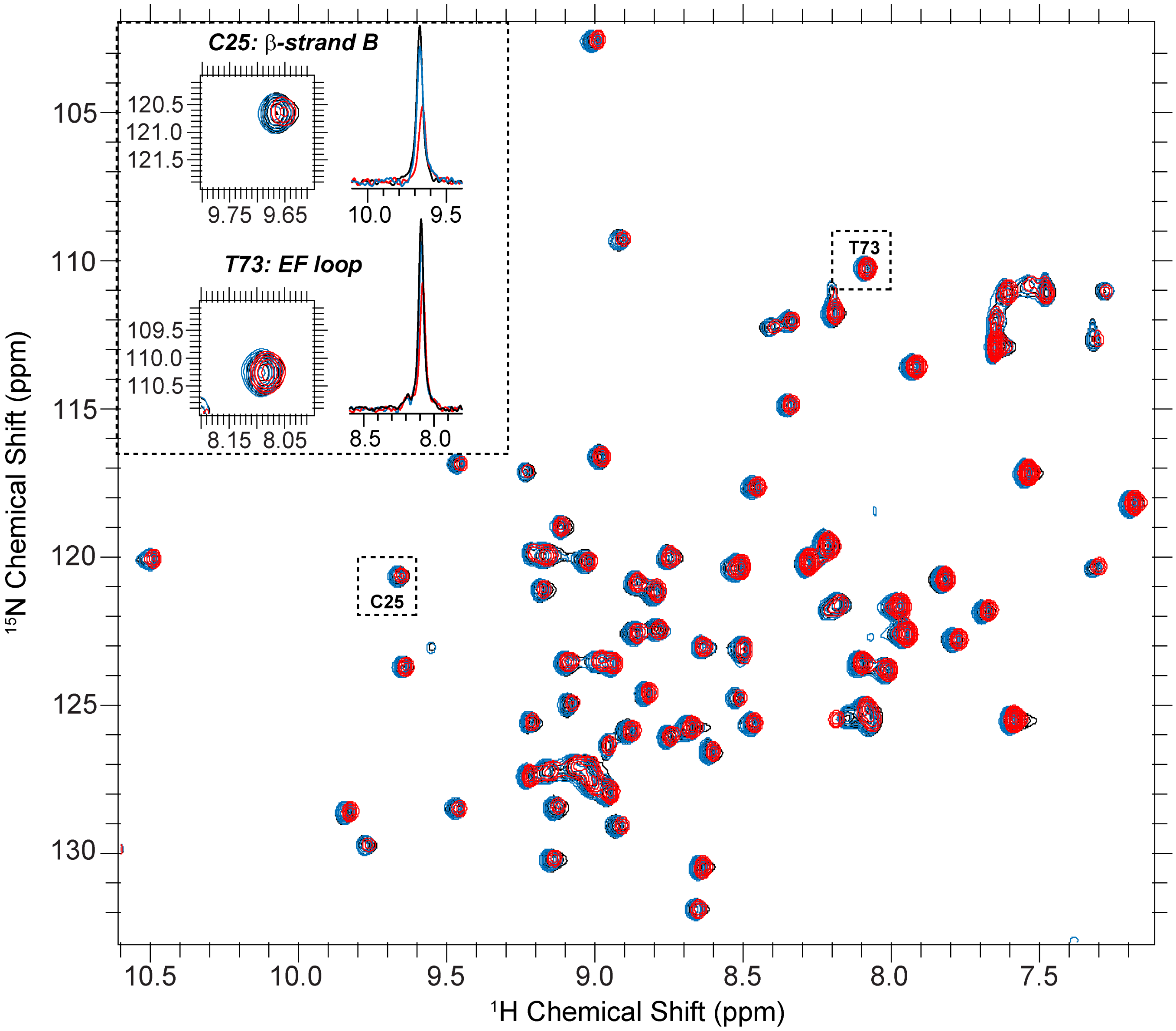


**Figure S1. Minimal chemical shift perturbation with residue-specific intensity losses observed by titration of collagen I into β$_2$m.** $^1$H–$^{15}$N-HSQC of 300 μM β$_2$m in TBS, pH 7.4 + 0.5 mg/ml casein in the absence (black) or presence of different concentrations of collagen I (blue- 0.12 mg/ml collagen I and red- 1.2 mg/ml collagen I). The inset shows a zoom-in on the 2D contours and the extracted $^1$H 1D projections of a residue that has a higher degree of peak intensity loss (Cys 25, $I/I_0$ = 0.48) and one that has a low level of intensity loss (Thr 73, $I/I_0$ = 0.76) upon addition of collagen I. Experiments were conducted in 10% D$_2$O at 700 MHz $^1$H Larmor frequency and 10$^o$C.

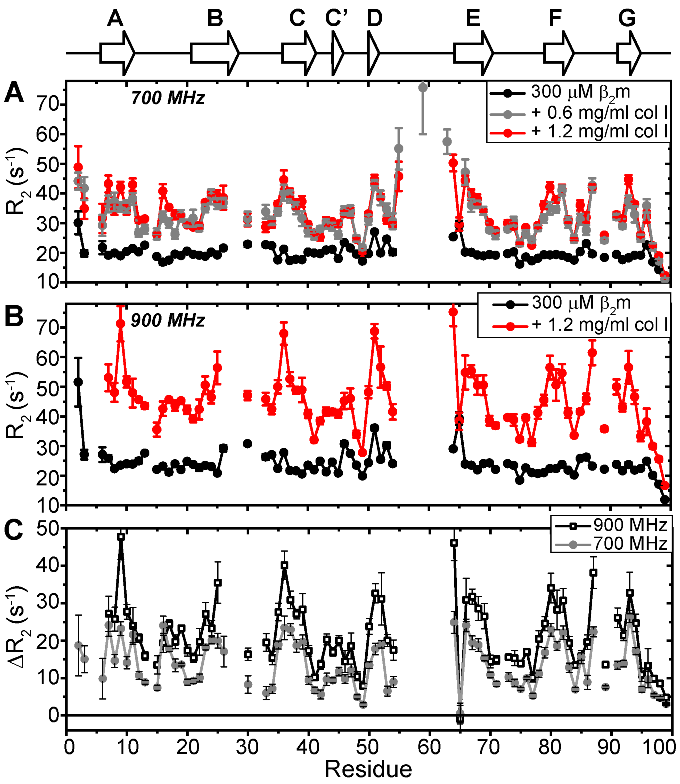


**Figure S2. Perturbation of $\beta_2$m $^{15}$N-$R_2$ at 900 MHz.** $^{15}$N-$R_2$ measurements of 300 µM $\beta_2$m in the absence (black) or presence (red) of 1.2 mg/ml collagen I in TBS, pH 7.4 + 0.5 mg/ml casein. Experiments were conducted in 10% $D_2O$ at 900 MHz $^1$H Larmor frequency and 10°C.

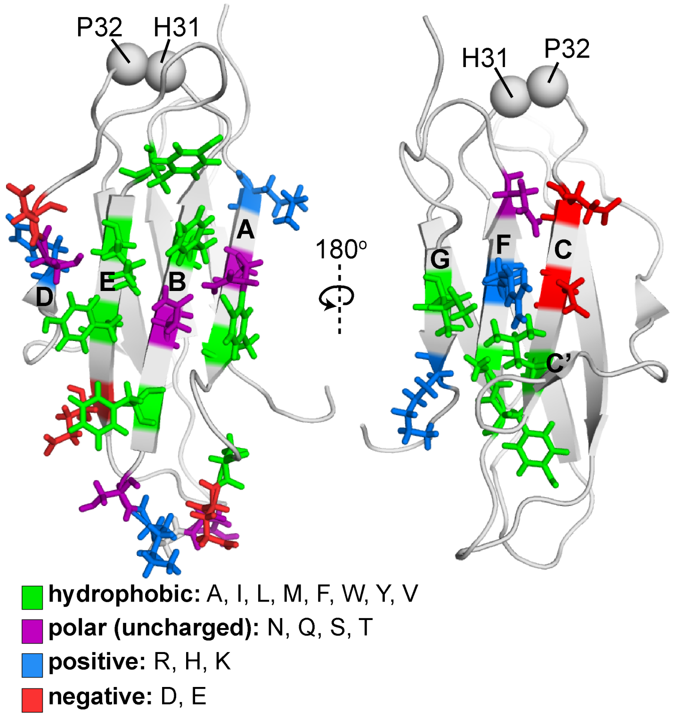


**Figure S3. Amino acid composition of the $\beta_2$m interface for collagen I interactions.** Amino acids determined to be at the $\beta_2$m–collagen I interface by $^{15}$N-DEST and that have side-chains oriented toward the interaction surface are shown in stick representation and colored by amino acid type (hydrophobic= green; polar, uncharged= purple; positive charge= blue; negative charge= red). His 31 and Pro 32 are shown as spheres. Both $\beta_2$m β-sheets are composed of a mixture of hydrophobic and hydrophilic amino acids, with the ABED β-sheet displaying several aromatic rings. Structural models are based on PDB: 2XKS[1].
